# Dynamics of activating and repressive histone modifications in *Drosophila* neural stem cell lineages and brain tumors

**DOI:** 10.1101/724039

**Authors:** Merve Deniz Abdusselamoglu, Lisa Landskron, Sarah K. Bowman, Elif Eroglu, Thomas Burkard, Robert E. Kingston, Juergen A. Knoblich

## Abstract

During central nervous system (CNS) development, spatiotemporal gene expression programs mediate specific lineage decisions to generate neuronal and glial cell types from neural stem cells (NSCs). However, little is known about the epigenetic landscape underlying these highly complex developmental events. Here, we perform ChIP-seq on distinct subtypes of *Drosophila* FACS-purified neural stem cells (NSCs) and their differentiated progeny to dissect the epigenetic changes accompanying the major lineage decisions *in vivo*. By analyzing active and repressive histone modifications, we show that stem cell identity genes are silenced during differentiation by loss of their activating marks and not via repressive histone modifications. Our analysis also uncovers a new set of genes specifically required for altering lineage patterns in type II neuroblasts, one of the two main *Drosophila* NSC identities. Finally, we demonstrate that this subtype specification in NBs, unlike NSC differentiation, requires Polycomb-group (PcG)-mediated repression.

**Summary statement:** Dynamic epigenetic landscape of Drosophila neural stem cell lineages.

## Introduction

During development of the central nervous system (CNS), neural stem cells (NSCs) divide asymmetrically to generate daughter cells with self-renewing capacity but also complex neurogenic and gliogenic lineages. Regulation of this process requires tightly and highly dynamic control of multiple cell fate decisions. For cells to commit to their ultimate cell identity, spatiotemporal gene expression programs are required. It is assumed that activation of lineage-specific genes and silencing of stem cell genes is accompanied by changes in chromatin states. How histone modifications change over time during neurogenesis *in vivo*, however, is not very well described.

The *Drosophila* larval CNS has become a key model for the fundamental mechanisms underlying brain development and chromatin states (Homem & Knoblich, 2012). The larval CNS is populated by distinct types of NSCs (called neuroblasts in *Drosophila*), which vary in abundance, neuronal output and division mode. Together, these NBs give rise to the majority of the adult brain’s neurons (Truman & Bate, 1988). The majority of the central brain NBs are of Type I (NBIs). Each NBI gives rise to another NBI and a ganglion mother cell (GMC), which divides once more to generate two differentiated neurons of glia. Type II neuroblasts (NBIIs) instead are a rare subpopulation with only 8 NBII per brain lobe (Fig. 1A) (Bello, Izergina, Caussinus, & Reichert, 2008; Boone & Doe, 2008; Homem & Knoblich, 2012; Sousa-Nunes, Cheng, & Gould, 2010). Unlike NBIs, NBIIs divide into one NBII and one transit-amplifying cell called intermediate neural progenitors (INPs). They generate many more neurons, because INPs continue to divide asymmetrically for 5-6 times, each time giving rise to a GMC that divides into two neurons or glia cells. (Bello et al., 2008; Boone & Doe, 2008; Homem & Knoblich, 2012). Other than lineage structure and size, cell markers can also be used to differentiate NB subtypes. While NBIs express both Asense (Ase) and Deadpan (Dpn) (Bowman et al., 2008), NBIIs only express Dpn (Bello et al., 2008). During neurogenesis, both NB subtypes divide asymmetrically to give rise to their respective progeny (Kang & Reichert, 2014). Brain tumors form if the asymmetric segregation of cell fate determinants during NB cell division is disrupted (Betschinger, Mechtler, & Knoblich, 2006; Juergen A Knoblich, 2010). Among these determinants are the TRIM-NHL protein Brain tumor (Brat) and the Notch inhibitor Numb (Arama, Dickman, Kimchie, Shearn, & Lev, 2000; Betschinger et al., 2006; Bowman et al., 2008; Jürgen A Knoblich, Jan, & Jan, 1995; Lee et al., 2006a; Lee, Wilkinson, Siegrist, Wharton, & Doe, 2006b). While Brat-depletion results in the generation of ectopic NBII-like tumor NBs (tNBs) at the expense of differentiated brain cells (Bowman et al., 2008), simultaneous loss of Brat and Numb causes the NBI-like tNBs to overproliferate (see Results).

**Figure 1.**
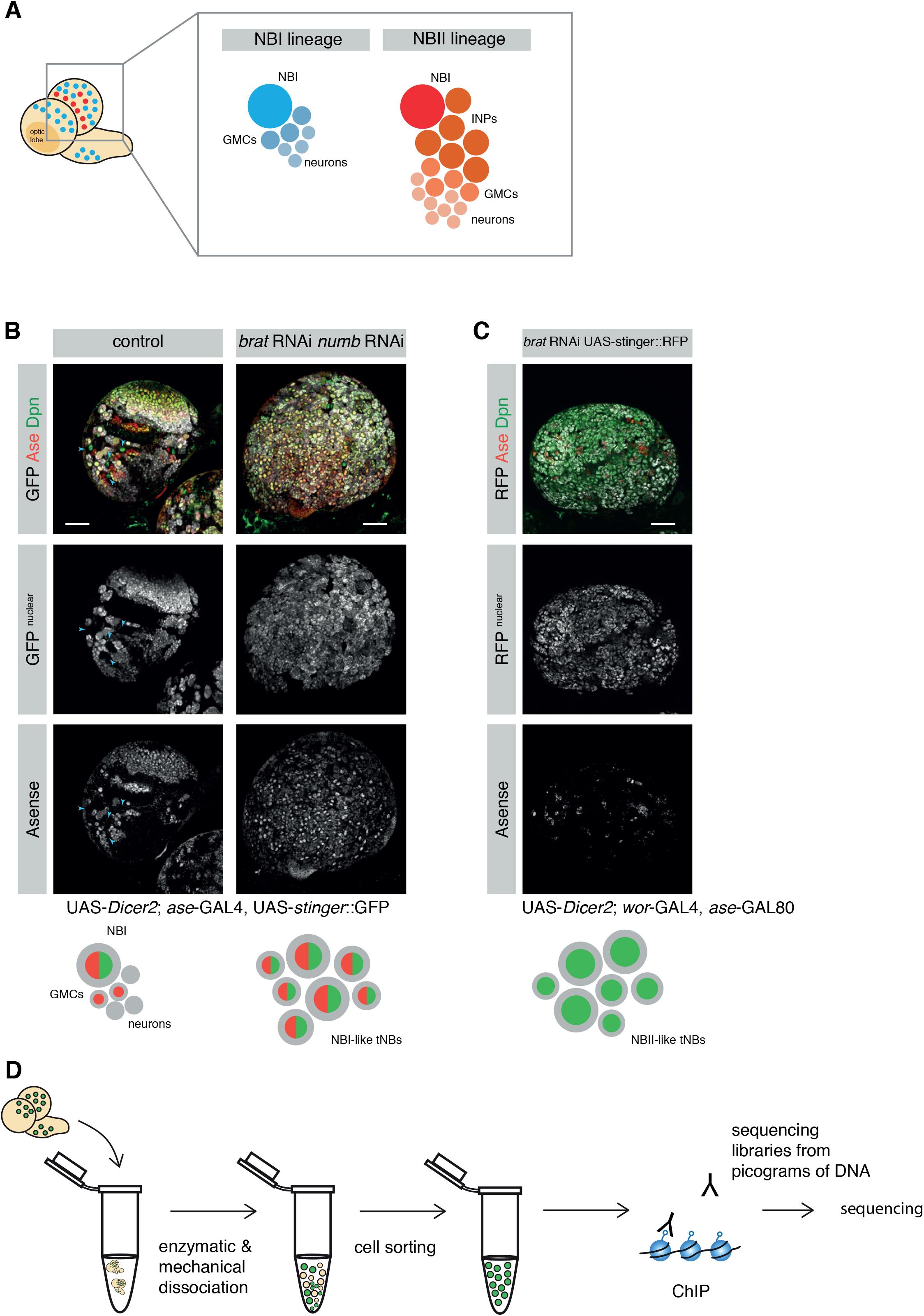
Strategy to investigate histone marks in specific NB lineages. (A) Cartoon depicting a larval brain with NBI (blue) and NBII (red) lineages. (A and B) Immunostainings with a scalebar = 50*μ*m. (B) The ase-GAL4 driver line marks NBI lineages with nuclear GFP but not NBII lineages (arrowheads). Combined knockdown of *brat* and *numb* results in ectopic Ase+ Dpn+ tNBs. Dashed line separates optic lobe (OL) and central brain. (C) *brat* depletion with the NBII specific driver line results in mainly Ase-Dpn+ tNBs. (D) Cartoon showing an overview of the ChIP-seq strategy.

In many cell types, transitions in chromatin states are regulated by the evolutionary conserved Polycomb (PcG) and Trithorax (TrxG) group proteins. PcG and TrxG have emerged as antagonistic regulators that silence or activate gene expression, respectively (Kingston & Tamkun, 2014; Levine et al., 2002; Schuettengruber, Chourrout, Vervoort, Leblanc, & Cavalli, 2007). These multimeric protein complexes regulate the transcriptional state of genes by post-translationally modifying amino acid residues of histone tails (Kingston & Tamkun, 2014; Levine, King, & Kingston, 2004). PcG proteins exert a repressive activity via two main complexes, the Polycomb repressive complexes 1 and 2 (PRC1 and PRC2). Although PRC1 and PRC2 can exist in various compositions and associate with context-specific accessory proteins, both PRC1 and PRC2 have been shown to contain a specific core set of proteins including subunits with catalytic activity (Bracken, Dietrich, Pasini, Hansen, & Helin, 2006; Simon & Kingston, 2009). Within PRC2, *Enhancer of zeste* (*E(z)* in *Drosophila*, EZH1/2 in mammals) catalyzes the trimethylation of lysine 27 on histone 3 (H3K27me3) (Cao & Zhang, 2004). H3K27me3 is recognized by PRC1, which in turn includes the histone 2A ubiquityltransferase *Sce* (RING1A/B in mammals) (de Napoles et al., 2004). Histone modifications associated with active transcription are deposited by TrxG proteins (Kassis, Kennison, & Tamkun, 2017), which counteract repressive histone acetylation or methylation marks, in particular by trimethylation of lysine 4 on histone 3 at active promoters (Byrd & Shearn, 2003; Dou et al., 2005; Petruk et al., 2001) (Kim et al., 2005).

Although well-known for their role in long-term transcriptional memory, PcG and TrxG complexes are highly dynamic during development and thus facilitate cellular plasticity (Kwong et al., 2008; Negre et al., 2006). In the last decade, it has been shown that PcG and TrxG complexes are crucial to ensure correct neurogenesis in mammals (Hirabayashi et al., 2009; Lim et al., 2009; Pereira et al., 2010) as well as in *Drosophila* (Bello, Holbro, & Reichert, 2007; Touma, Weckerle, & Cleary, 2012). Despite the strength of genetic *in vivo* experiments, however, global analysis of the histone modifications underlying their function, and therefore target genes, has mainly been performed *in vitro*. This constitutes a real knowledge-gap as recent studies demonstrated that the chromatin states may vary significantly between *in vivo* tissues and their related *in vitro* cell lines, mainly due to culture conditions (R. Xie et al., 2013; Zhu et al., 2013). Given also that epigenetic changes are highly context – and developmental time-dependent, providing *in vivo* datasets to investigate chromatin states of different cell types in complex tissues will increase our understanding of how the epigenetic landscape dynamically defines cellular states.

In recent years, *in vivo* studies made use of *Drosophila* to shed light on the dynamics of chromatin state changes during embryonic neural differentiation (Ye et al., 2016) and during larval stages (Aughey, Estacio-Gómez, Thomson, Yin, & Southall, 2018; Marshall & Brand, 2017). Profiling the binding of chromatin remodelers has highlighted the plasticity of chromatin states during differentiation (Marshall & Brand, 2017). Although binding of chromatin factors is associated with active or repressive chromatin, binding does not necessarily reflect downstream histone modifications. For example, the histone marks can change drastically between parasegments of the *Drosophila* embryo while the occupancy of PcG proteins remains unchanged (Bowman et al., 2014). Thus, investigating the dynamics of chromatin states based on chromatin marks is crucial for understanding the functional specialization of cells during development. Moreover, how PcG/TrxG complexes target genes on the chromatin level between different subtypes of progenitor cells during neuronal differentiation, or tumorigenic transformation has remained elusive.

Here, we use the *Drosophila* larval CNS to track *in vivo* changes of histone modifications not only upon differentiation, but also between different populations of neural stem cells and their tumorigenic counterparts. We developed a FACS-based method to sort different cell types and perform ChIP-Seq for the active histone mark, H3K4me3, and the repressive mark, H3K27me3. Our FACS-based approach provides an in vivo dataset that reveals dynamic histone modifications during neuronal differentiation. In particular, we observed that self-renewal and cell division genes are repressed independently of H3K27me3 levels. In contrast, we further show that H3K27me3-mediated repression is crucial for silencing lineage-specific stem cell factors, including known factors as wells as a new set of genes that are specific to NBIIs. Finally, we present genetic evidence for the requirement of these new NBII-specific factors for self-renewal and demonstrate the role of PcG complexes in defining different subtypes of neural stem cells.

## Results

### Profiling repressive and active histone modifications of neural stem cells and neurons

H3K4me3 and H3K27me3 are two major histone modifications associated with TrxG-activated and PcG-repressed states, respectively. However, these histone modifications have not yet been analyzed independently in distinct subtypes of neural stem cells in *Drosophila*. To analyze H3K4me3 and H3K27me3 histone marks in different brain cell types by ChIP-Seq, we combined genetic labeling with a protocol for generating sequencing libraries from picogram quantities of DNA (Bowman et al., 2014). The NB subtype-specific GAL4 drivers *ase*-GAL4 (NBI lineage-specific) and *wor*-GAL4, *ase*-GAL80 (NBII lineage-specific), allowed us to preferentially label distinct NB lineages with nuclear-localized fluorophores (stinger::GFP or RFP). Indeed, GFP expressed by *ase*-GAL4 exclusively labeled NBI lineages (positive for both Dpn and Ase) and was not expressed in NBIIs (only Dpn+) (Fig. 1B). To amplify the production of rare NBIIs and at the same time generate tumor NBs (tNBs), RNAi constructs against the cell fate determinants *brat* and *numb*, were expressed using the mentioned driver lines. Depletion of *brat* in larval brains with a NBII-specific driver resulted in the overproliferation of NBII-like tNBs, evident by an increase in Dpn+, Ase-cells (Fig. 1C). In contrast, simultaneously depleting *numb* and *brat* by an NBI-specific driver resulted in overgrowth of (Dpn+, Ase+) NBI-like tNBs (Fig. 1B). Therefore, this strategy allowed us to generate fluorescently labeled distinct NB cell types and neurons.

Besides central brain NB lineages, the larval brain consists of embryonic neurons, mushroom body neuroblasts and cells of the optic lobes. To avoid impurities from these structures NBI, neurons and tNBs were isolated according to fluorescence intensity and cell size by flow cytometry (Berger et al., 2012; Harzer, Berger, Conder, Schmauss, & Knoblich, 2013). Purified cell populations were then analyzed by ChIP-seq for H3K27me3 and H3K4me3 histone modifications (Fig. 1D).

The H3K4me3 signal peaked around transcriptional start sites (TSSs), whereas the H3K27me3 signal occurred in broad domains covering gene bodies. For example, in all cell types the ubiquitously expressed gene *RNA polymerase II subunit 215kDa* contained a H3K4me3 peak at the TSS which was devoid of H3K27me3 signal (Fig. S1A). In contrast, the gene *caudal* showed no H3K4me3 peak, but instead high H3K27me3 levels over the gene body (Fig. S1B). This is in accordance with the fact that the function of *caudal* is mostly restricted to the larval digestive system. Moreover, *caudal* is not expressed in the larval CNS (modENCODE data and (Berger et al., 2012)) and has been shown to inhibit neuroblast specification upon misexpression (Birkholz, Vef, Rogulja-Ortmann, Berger, & Technau, 2013).

### Self-renewal and cell cycle genes are repressed during differentiation in a H3K27me3-independent manner

To investigate changes in the epigenetic landscape during neurogenesis, we collected NBIs and neurons as described above in duplicates. We subtracted the individual inputs from their respective samples to generate coverage tracks. The read counts for H3K27me3 localization were analyzed over the whole gene body, but the reads for H3K4me3 were counted 500 bp downstream of the TSS. From this data, regions with differential signals were identified between different cell types. Finally, we performed unsupervised hierarchical clustering analysis on differentially marked genes in NBIs and neurons (Fig. 2A) identifying five distinct groups of genes. Three clusters were dependent on H3K27me3-mediated repression. Cluster 1 showed a decreased H3K27me3 signal upon differentiation, while cluster 3 showed an increased H3K27me3 signal in neurons. These clusters were not enriched for genes of a particular pathway or biological process when analyzed with gene ontology enrichment analysis. Another example of H3K27me3-mediated repression was cluster 5. While cluster 1 and 3 showed changes in H3K27 signal and no or mild changes in H3K4me3, genes in cluster 5 (22 genes) showed a drastic switch from a H3K4me3+ H3K27me3-to a H3K4me3-H3K27me3+ chromatin state upon differentiation. Cluster 5 included the lncRNAs *cherub*, *pncR002:3R* and *sphinx* and transcription factors *nab* and *vvl* (Fig. 2B).

**Figure 2.**
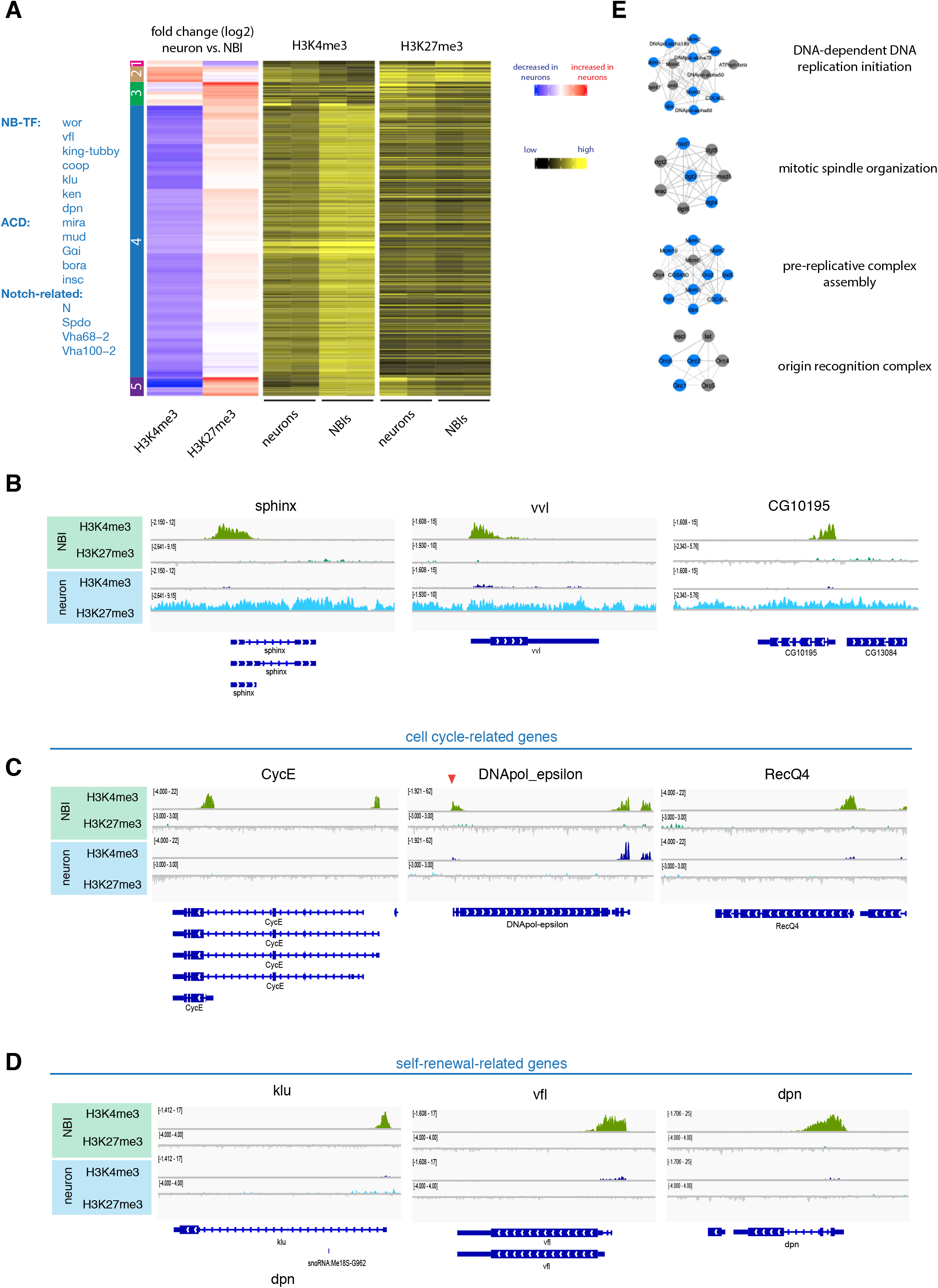
Changes of active and repressive histone modifications upon differentiation. (A) Unsupervised hierarchical clustering analysis of gene log2 foldchange between NBIs and neurons. NB-related genes of cluster 4 according to literature are indicated blue. (B) ChIP-seq tracks of representative examples for genes of cluster 5. ChIP-seq tracks of representative examples for cell-cycle-related genes (C) and self-renewal-related genes (D). (E) Examples of mitosis related protein complexes. Blue indicates genes found in cluster 4.

The other two clusters (Cluster 2 and 4) were mainly dependent on changes in H3K4me3 signal. Cluster 2 showed an increase in H3K4me3 levels in neurons, while cluster 4 contained a large number of 318 genes and was characterized by a loss of H3K4me3 upon differentiation. These gene loci had either a small increase in H3K27me3 or were completely devoid of both marks (H3K4me3 and H3K27me3) in neurons (Fig. 2A, C, D). Gene ontology enrichment analysis (Supplement table 1) showed that cluster 4 genes were enriched for genes involved in self-renewal (e.g. stem cell proliferation p=0,002) and mitosis-related processes (e.g. DNA replication p=3,04E-21), which are both processes that cease upon differentiation. In support of this finding, protein complexes essential for cell division were enriched in cluster 4 (Fig. 2E), whereas *bona fide* NB self-renewal transcription factors such as *vfl*, *klu*, and *dpn* as well as asymmetric cell division regulators (*Gαi, mud, insc, bora*) appeared in cluster 5. This suggests that during differentiation stem cell-promoting genes lack a H3K27me3 mark and that their repression could be mediated through mechanisms independent of PcG. This result is corroborated by previous data indicating that the genes *mira*, *CycE*, *stg* and *dpn* are enriched in HP1-associated chromatin in *Drosophila* neurons (Marshall & Brand, 2017).

Thus, these data suggest that a small group of genes is controlled by H3K27me3-repression upon differentiation, whereas most stem cell–related genes are turned off via an additional mechanism, potentially involving HP1 enrichment. This result is surprising considering H3K27me3 datasets indicate spreading of PcG-repressed regions upon neural differentiation in mammals (Södersten et al., 2018; Zhu et al., 2013) and suggests that different strategies of epigenetic control of neurogenesis have been established across evolution.

### Subtype-specific neuroblast genes are controlled by TrxG and PcG

Next, we wanted to address whether alterations of histone modifications can be observed between different types of NB lineages. To this end, we made use of tNBs which are of a different origin. RNAi of both *brat* and *numb* induced tumors made of NBIs while the depletion *brat* alone initiates tumors consisting of NBIIs (Fig. 1B-C). We reasoned that features occurring in NBII-like tNBs but absent in NBI-like tNBs would likely be specific to NBII lineages rather than due to tumorigenesis. We performed hierarchical clustering analysis between NBIs, NBI-like tNBs and NBII-like tNBs as described above and identified two NBII-specific sets of genes (Fig. 3A and Fig. S2). The first cluster of genes showed a decrease in H3K4me3 in NBII-like tNBIIs compared to NBIs and no or only mild changes when compared to NBI-like tNBIs (NBII cluster1). Moreover, these loci showed no or only modest increases in H3K27me3. As a key example, among this set of genes, we found the NBI-specific transcription factor *asense*, which showed clear H3K4me3 signals in NBI and *brat numb* depleted NBI-like tNBs but no signal in NBII-like tNBs (Fig. S3A).

**Figure 3.**
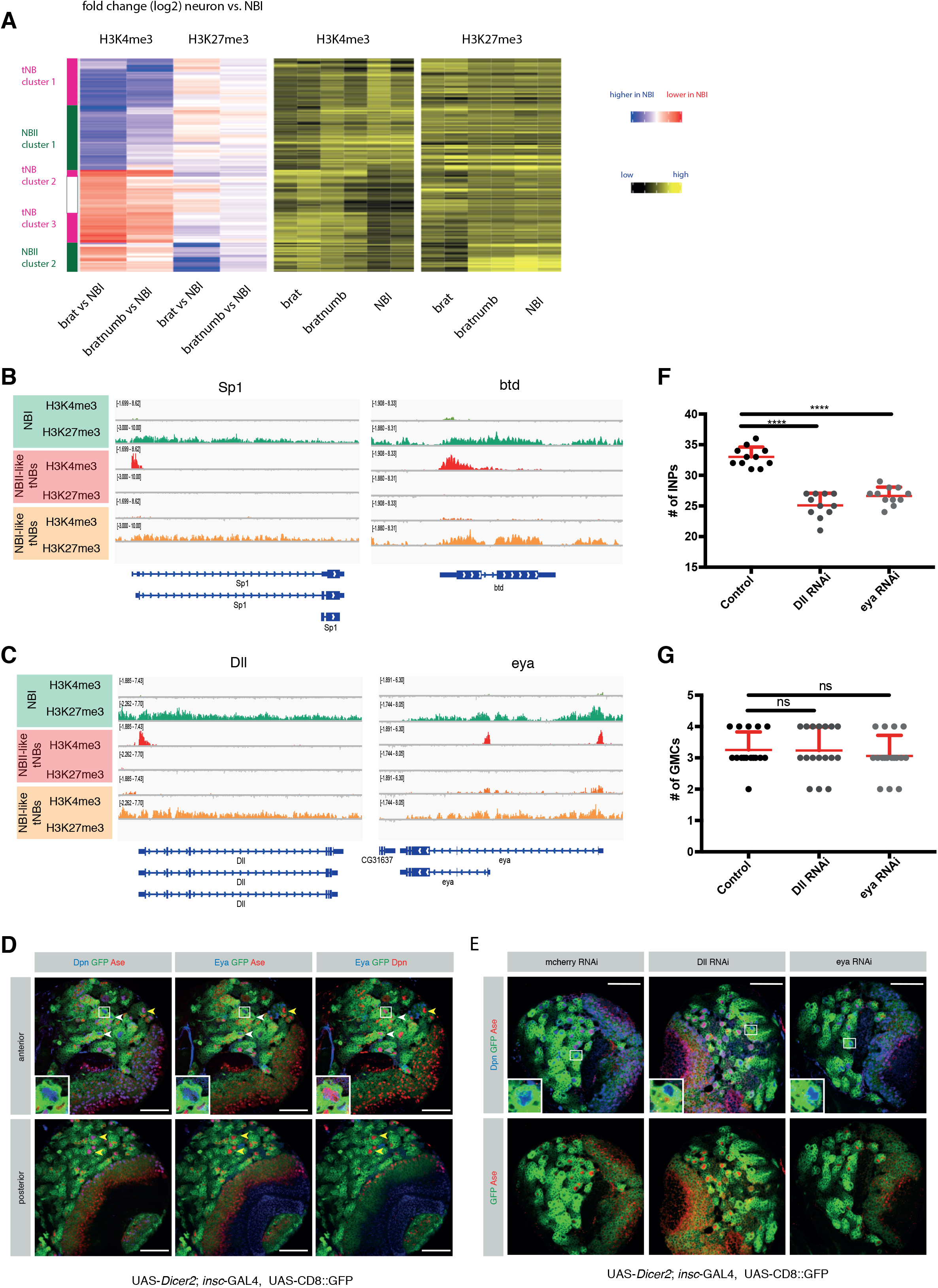
Comparison of different NB subtypes identifies PcG and TrxG-dependent NBII-specific factors. (A) Unsupervised hierarchical clustering analysis of gene log2 foldchange between NBI, NBI-like tNB and NBII-like tNB (relevant section is shown, for full heatmap see Fig. S5.). ChIP-seq tracks of the known NBII factors *Sp1* and *btd* (B) and novel NBII-specific factors *Dll* and *eya* (C). (D) Eya immunostaining in type I and type II neuroblasts. Scale bar 50 *μ*m. Driver line used was UAS-*dicer2*; *insc*-Gal4, UAS-CD8::GFP. (E) Immunostainings of larval brains expressing RNAi against of Dll or eya show smaller NBIIs (blow-up). Scale bar is 50 *μ*m. Quantification of immediate progenies of NBIIs (F) and NBIs (G). Driver line used was UAS-*dicer2*; *insc*-Gal4, UAS-CD8::GFP. Mean ± SD is shown. (for INPs (F) n=11, control = 33±1.61, *Dll* = 25.09±1.97 and *eya* = 26.64±1.43, and for GMCs (G) control = 3.25±0.57 (n=16), *Dll* = 3.23± 0.75 (n=17) and *eya* = 3.05±0.65 (n=17)). One-way ANOVA test was used and ****p < 0.0001, ns = not significant. n numbers are lineages quantified.

In the second gene cluster (NBII cluster 2), NBII-like tNBIIs showed increased H3K4me3 signal and lower H3K27me3 occupancy. Interestingly, previous genetic evidence has suggested a role of *trithorax* in maintaining different subtypes of neuroblast lineages. In particular, the two loci *buttonhead* and *Sp1* are required to specify NBII from neuroectoderm (Álvarez & Díaz-Benjumea, 2018) and to maintain NBII lineages (Komori, Xiao, Janssens, Dou, & Lee, 2014; Y. Xie et al., 2014). Indeed, both genes are H3K4me3 positive in NBII-like tNBIIs and have reduced intensity of H3K27me3 signal in NBI-derived numb tumors as well as NBIs (Fig. 3B). Our clustering identified additional genes with a similar pattern (Fig. 3A). Two of these H3K4me3 positive NBII-like NBII-specific genes were the homeodomain transcription factor Distal-less (Dll) and the transcriptional coactivator eyes absent (eya). We decided to further focus on and characterize the contribution of these two genes in NBII specification as previous work showed that Dll enhancer was active in NBII lineages (Izergina, Balmer, Bello, & Reichert, 2009) and eya was mainly expressed in NBIIs (which we confirmed by immunostaining (Fig. 3D)). Dll-or eya-depleted brains resulted in smaller NBII cells (Fig. 3E and Fig. S2B) with a reduced number of INPs (Fig. 3F), which indicates reduced stemness (Song & Lu, 2011; Wissel et al., 2018). In contrast, NBI lineages showed normal NB growth and unaffected GMC numbers (Fig. 3G and Fig. S2C). Thus, our data indicate that these two genes are required to maintain NBII lineages. This would further suggest that NB subtype-specific genes are regulated by PcG and TrxG.

Finally, tNB-specific changes were mostly H3K4me3 changes (reduction in tNB cluster 1 and gain in tNB cluster 2+3) (Fig. 3A and Fig. S2) while only minor changes were observed for H3K27me3. This suggests that tumor specific changes are mediated by TrxG proteins rather than PcG.

### PRC 1 and 2 are required for neuroblast maintenance

Our data suggest that both TrxG and PcG complexes play an important role in maintaining NBI and NBII identities. TrxG-dependent maintenance of NBII identity was shown to rely on the target genes *buttonhead* and *Sp1* (Álvarez & Díaz-Benjumea, 2018; Komori et al., 2014). By contrast, the role of PcG in NB-subtype specification, besides HOX gene repression, remains largely unexplored. During brain development, loss of PcG repression leads to ectopic expression of HOX genes, which in turn induces apoptosis and depletion of both type I and type II NBs (Bello et al., 2007). In accordance with these previous findings, our ChIP-seq data revealed high levels of H3K27me3 at the two HOX gene clusters Antennapedia and Bithorax complex in both NBIs and NBIIs (Fig. S4A).

To investigate whether PRC-mediated repression is only required to prevent HOX gene-induced apoptosis or plays a broader role in neurogenesis, we blocked apoptosis by expressing the baculovirus caspase inhibitor gene P35. To this end, RNAi constructs against components of PRC2 (*E(z), Su(z)12*) or PRC1 (*Sce*) were expressed in NBs using the general NB driver line *insc*-GAL4, which resulted in a great decrease in NBI and NBII cell numbers (Fig. 4A). Upon p35 expression in a PcG-depleted background, GFP+ NB lineages could be restored (Fig. 4B) as previously reported (Bello et al., 2007).

**Figure 4.**
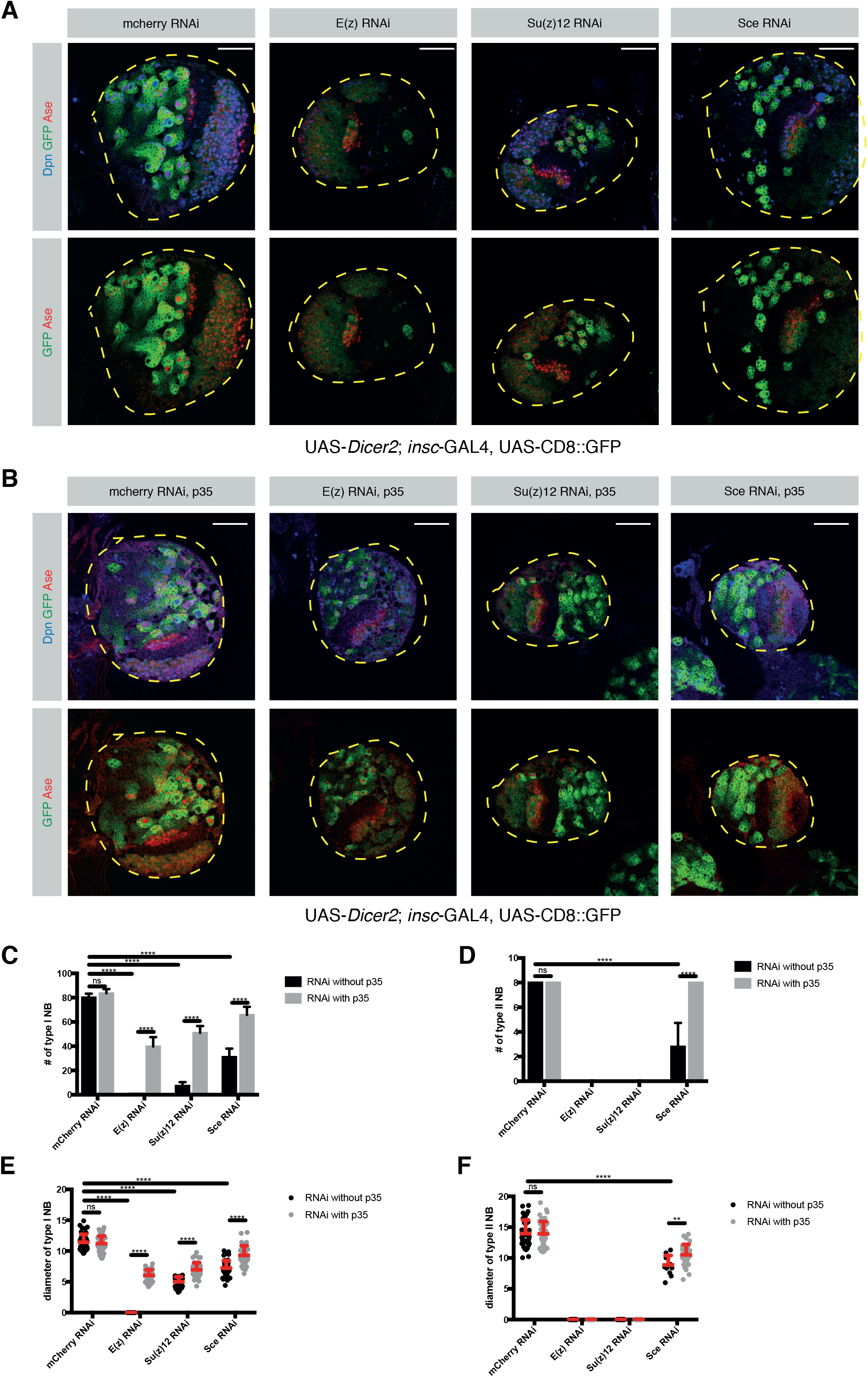
PRC 1 and 2 are required for NB maintenance. (A) Loss of PRC1 and 2 causes a significant decrease in NB numbers. Larval brain lobes expressing RNAi against mCherry, *E(z), Su(z)12* and *Sce.* Lobes are outlined in yellow dashed lines. Scale bar 50 *μ*m. Driver line used was UAS-*dicer2*; *insc*-Gal4, UAS-CD8::GFP. (B) Larval brains expressing apoptosis inhibitor P35 together with PRC RNAi constructs in (A). Lobes are outlined in yellow dashed lines. Scale bar 50 *μ*m. Driver line used was UAS-*dicer2*; *insc*-Gal4, UAS-CD8::GFP. (C) Quantification of NBI numbers in mCherry, E(z), Su(z)12, and Sce-depleted larval brains with and without P35 expression. n= 5 brain lobes. Mean ± SD (mCherry = 80.2±3.11, mCherry+P35 = 83.6±3.46, *E(z)* = 0.2±0.44, *E(z)*+P35 = 39.8±7.66, *Su(z)12* = 7.4±2.96, *Su(z)12*+P35 = 51±5.61, *Sce* = 31.2±6.83 and *Sce*+P35 = 65.75±6.7). Two-way ANOVA test was used and ****p < 0.0001. (D) Quantification of NBI diameter in mCherry, E(z), Su(z)12, and Sce-depleted larval brains with and without P35 expression. Mean ± SD (mCherry = 10.64±1.6 (n=50), mCherry+P35 = 11.61±1.93 (n=50), *E(z)* = NA, *E(z)*+P35 = 6.05±0.9 (n=50), *Su(z)12* = 6.01±1.14 (n=34), *Su(z)12*+P35 = 6.76±1.3 (n=50), *Sce* = 7.21±1.22 (n=50), and *Sce*+P35 = 9.31±1.54 (n=50)). Two-way ANOVA test was used and ****p < 0.0001. n = NBI numbers quantified. (E) Quantification of NBII numbers in mCherry, E(z), Su(z)12, and Sce-depleted larval brains with and without P35 expression. n= 5 brain lobes. Mean ± SD (mCherry = 8, mCherry+P35 = 8, *E(z)* = NA, *E(z)*+P35 = NA, *Su(z)12* = NA, *Su(z)12*+P35 = NA, *Sce* = 2.8±1.92 and *Sce*+P35 = 8). Two-way ANOVA test was used and ****p < 0.0001. (F) Quantification of NBII diameter in mCherry, E(z), Su(z)12, and Sce-depleted larval brains with and without P35 expression. Mean ± SD (mCherry = 13.96±2.19 (n=40), mCherry+P35 = 13.91±2 (n=40), *E(z)* = NA, *E(z)*+P35 = NA, *Su(z)12* = NA, *Su(z)12*+P35 = NA, *Sce* = 8.85±1.56 (n=12) and *Sce*+P35 = 10.5±1.7 (n=32)). Two-way ANOVA test was used and ****p < 0.0001. n = NBII numbers quantified.

Although RNAi constructs were expressed with the same driver line in NBIs and NBIIs, blocking apoptosis in PRC2-depleted lineages restored NBI but not NBII cell numbers (Fig. 4C, D). In contrast, the number of NBs in PRC1-depleted brains was restored, suggesting PRC1, unlike PRC2, seem to only target the HOX genes and therefore prevent apoptosis of NBs (Fig. 4D). However, these restored NBIIs still exhibited a smaller cell size (Fig. 4F) than their control NBIIs. These results indicate that in addition to its function as anti-apoptotic in both type I and type II NB, that NBIIs require PRC2 is required specifically in NBIIs to maintain self-renewal potential. These data therefore suggest that PcG-dependent repression targets more genes in addition to the HOX genes to maintain NBII.

### PcG proteins prevent premature NB differentiation

Although the number of NBIs was restored in apoptosis-inhibited PcG-depleted conditions, the NBI cell size was reduced (Fig. 4E). NBs must maintain a certain growth rate to maintain their self-renewal potential and prevent differentiation (Song & Lu, 2011). We therefore analyzed these apoptosis-inhibited NBIs and their self-renewal potential. RNAi constructs against PRC1 and PRC2 components were expressed together with P35, and NBs were analyzed at 6h after pupal formation (APF), timepoint at which NB start to exit proliferation (Fig. S4B). While the number of NBI in both PRC1- and PRC2-depleted brains were restored in third instar larval brains upon P35 expression, NB numbers were significantly decreased at 6h APF along with the fact that the diameter of the remaining ones was significantly lower compared to control (Fig. S4C, D). Altogether, these data show that even though the number of NBs were restored in apoptosis-inhibited, PRC-depleted lineages, these NBs fail to maintain their self-renewal potential as reported by their smaller size and early differentiation compared to their wild-type counterparts. These results altogether indicate that PcG proteins are required to maintain stemness both in type I and type II NBs but with different sensitivities.

To address the physiological consequences of premature NB differentiation, we analyzed the viability of PcG-depleted flies. RNAi-mediated knockdown of PcG proteins with and without p35 expression using *insc*-GAL4, led to lethality during development (data not shown). This observation further confirms our previous results that neurogenesis of NBI lineages is not fully restored. However, this approach suffers from the caveat that the *insc* promotor is active in some cells of the larval gut and salivary glands. To exclude that lethality could originate from abnormal development of other tissues, we next used a brain-restricted NBI lineage-specific driver line *ase*-GAL4. Similar to insc-Gal4, ase-Gal4-mediated loss of PcG proteins led to a decline in NBI numbers and size, which could be rescued by blocking apoptosis (Fig. S5A, B). These phenotypes were nonetheless weaker, which we could explain by the strength of *insc-*Gal4, is higher expressed than *ase* in NBIs (Berger et al., 2012). However, immunostainings could confirm that PcG RNAi upon p35 rescue NBI lineages using *ase*-GAL4 still showed a significant decrease in H3K27me3 signal (Fig. S5C). Therefore, these results further confirm that PcG promotes self-renewal beyond preventing apoptosis also in NBI.

When PcG proteins were depleted with *ase*-GAL4 driven RNAi during NBI development, the majority of eggs failed to develop into adult flies (Fig. S5D). Between 3-18% of laid eggs hatched, but flies showed neurological abnormalities, and became stuck in the fly food leading to death. Similarly, preventing apoptosis in these PcG knockdown backgrounds did not rescue the number of viable flies (Fig. S5D). Therefore, NBIs and NBIIs depend on PcG proteins for proper neuron production, although at different sensitivities. In summary, these results suggest that PRC1 and 2 maintain NB neurogenesis by silencing genes that induce apoptosis and genes whose expression leads to differentiation.

## Discussion

We provide a resource of histone modification datasets for different types of neural stem cells and their differentiated progeny. In combination with chromatin accessibility (Aughey et al., 2018) and binding maps of chromatin remodelers (Marshall & Brand, 2017) of *Drosophila* brain cells, we hope that our dataset will serve as an useful community resource. We show that during differentiation, stem cell identity genes are silenced in a PcG-independent manner, which supports previous findings showing that these genes are silenced through HP1 enriched chromatin (Marshall & Brand, 2017). Additionally, PcG-mediated silencing is unlikely to instruct the stepwise inactivation of stem cell genes during differentiation as loss of H3K27me3 did not induce ectopic NBs.

Here, we take advantage of *in vivo* genetic labeling to investigate chromatin dynamics of different NB subtypes. As the type II NBs are very lowly abundant, we used tumor NBs of type I and type II origins as a proxy in order to obtain enough material to be able to compare these two cell types. We further validated each change observed by comparing tumor to healthy type I NBs and excluded artifacts due to the tumorigenic state of the cells. Our data show that both TrxG and PcG are required to establish NBII identity. We identify a set of NBII-specific genes, including previously identified *btd* (Komori et al., 2014) and *Sp1* (Álvarez & Díaz-Benjumea, 2018). We further identified *Dll* and *eya* which are specifically required for NBII maintenance. It has been previously described that *btd* acts as an activator of *Dll* in the development of the ventral imaginal discs (Estella, Rieckhof, Calleja, & Morata, 2003). This suggests that in NBII-identity specification the Trithorax-target *btd* could act together with *Dll* and *eya*. Such a mechanism would explain why the loss of *btd* causes a distinct phenotype compared to the loss of *Dll* and *eya*. Interestingly, a NBI to NBII conversion is observed only in 18% of NBIs ectopically over-expressing *btd* indicating that either cofactors are missing or that the chromatin of *btd* targets is inaccessible (Komori et al., 2014). Our data of NB subtype-specific genes being characterized by H3K27me3 repressive chromatin favor the latter.

Therefore, as opposed to TrxG-activated stem cell and mitosis genes, the repression of NBII-specific genes is ensured by PcG-mediated H3K27me3 histone modifications suggesting that Polycomb plays a role in defining the diversity of neural stem cell lineages. Moreover, our data indicates that PcG repression is required not only for the silencing of HOX genes but also for the self-renewal capacity of NBs. Unlike TrxG (Komori et al., 2014), the loss of catalytic subunits of PcG complexes did not convert NBIIs to NBIs or vice versa. This suggests that NB subtype-specification cannot be explained solely by an absence of repression but requires a further activation mechanism. Strikingly, loss of PcG complexes caused a significant decrease in the numbers of NBs. Interestingly, across all the cell types, developmental genes such as *cad, eve, peb, scr*, and *slp1*, as well as genes involved in embryonic NB temporal patterning (*hb, kr, pdm, cas* and *grh*), are heavily marked with H3K27me3. It is therefore possible that PcG-mediated repression is required to silence these developmentally crucial genes in addition to the HOX genes. Thus, the observed reduction in NB stemness might be caused by the de-repression of these genes.

Besides an overall decreased NB maintenance, we observed increased sensitivity of NBII lineages upon reduction of PRC2 activity. Interestingly, *opa* and *ham*, two previously described temporal switch genes in NBII lineages (Abdusselamoglu, Eroglu, Burkard, & Knoblich, 2019), are also enriched with H3K27me3 in NBs. Ectopic expression of these genes limits self-renewal of NBs and causes NBs to disappear (Abdusselamoglu et al., 2019; Eroglu et al., 2014). In the future, investigating the downstream targets of PcG in different NB subtypes could reveal the underlying mechanisms of subtype-specification. In conclusion, our data provide a useful resource to investigate how chromatin state dynamics orchestrate the diversity and correct progression of neural stem cell lineages.

## Materials and Methods

### Fly strains

UAS-*Su(z)12* RNAi (BL 31191), UAS-*E(z)* RNAi (BL36068), UAS-*Sce* RNAi (VDRC 106328), UAS-p35 (BL5072, BL5073, both were tested for functionality), UAS-Dll RNAi (VDRC 101750), UAS-eya RNAi (VDRC 108071).

### Immunofluorescence

Brains were dissected and fixed for 20 min in PBS with 5% PFA with 0.1% TritonX-100. After three washes with 1XPBS with 0.1% TritonX-100 (PBST), brains were incubated for 1 hour in blocking solution (PBST with 3% Normal goat serum), incubated with blockings solution with primary antibodies and washed again three times with PBST. Secondary antibodies (1:500, goat Alexa Fluor®, Invitrogen) were added for one to two hours and then removed with three PBST washes. Brains were mounted in Vectashield Antifade Mounting Medium (Vector Labs). Primary antibodies used were: rat anti-Asense (1:500, (Eroglu et al., 2014)), guinea pig anti-Deadpan (1:1000, (Eroglu et al., 2014)), H3K27me3 (1:500, Active Motif 39155).

### Microscopy

Images were recorded on Zeiss Confocal 780. Images of different conditions in one panel were recorded using the same settings.

### Isolation of NBs using FACS

NB-sized cells were sorted from third instar larval brains according to GFP/RFP signal and cell size as previous described (Berger et al., 2012; Harzer et al., 2013). Briefly, brains were collected in 1X Rinaldini solution and then enzymatically and mechanically dissociated in Schneider’s medium supplemented with FBS (10%), PenStrep (2%), Insulin (20 *μ*l/ml), L-Glutamine (20mM), L-Glutathione (40 *μ*g/ml), 20-Hydroxyecdyson (5 *μ*g/ml).

### Statistics

Statistical analyses were performed with GraphPad Prism 7. Experiments were not randomized. Sample sizes were estimated depending on the previous experiences with similar setups and the investigator wasn’t blinded. Two-way ANOVA was used to assess statistical significance between multiple samples, while unpaired two-tailed Student’s *t*-test was used between two samples.

### ChIP-Seq

#### Preparation of soluble chromatin

50000 sorted cells of interest were pelleted by centrifuging at 3000 rpm for 10 min. The cell pellet was resuspended in complete media. Fixation was performed with formaldehyde (final concentration 1%) for 5 min at room temperature. After quenching with glycine (final concentration 125 mM) for 3 min at room temperature, cells were centrifuged at 3000 rpm for 10 min and the supernatant was discarded. Cells then were resuspended in 100 *μ*l 1X PBS with CaCl_2_ (final concentration: 1mM) and TritonX100 (final concentration: 0.1%) and incubated with 5 Units micrococcal nuclease (Worthington Biochemical, LS004798) at 37°C for 3 min. After incubation, the sample was immediately transferred on ice and 2.5 *μ*l 0.5 M EDTA, 6.25 *μ*l 0.2 M EGTA and 1.25 *μ*l 1X PBS were added in order to stop the reaction. After adjusting the sample volume to 300 *μ*l with 1X PBS and sonication was performed with a microtip sonicator (OmniZRuptor 250, Omni International; microtip, power output: 20) for 20 seconds in a prechilled metal. Once sonication was over, the sample was snap frozen in liquid nitrogen. Fragment size was assessed using the Agilent High Sensitivity DNA Assay.

#### Chromatin Immunoprecipitation

The volume of thawed chromatin samples was adjusted to 500 *μ*l with 50 *μ*l 10X lysis buffer (1 mM EDTA, 10 mM Tris-HCl pH8, 0.1% Na-Deoxycholate, 0.5% N-lauroylsarcosine, 1% TritonX100), 140 *μ*l water and 10 *μ*l of 50X complete protease inhibitor. After 5min incubation on ice, samples were spun down at maximum speed for 10 min at 4°C. While supernatant was transferred to fresh tubes, 5 *μ*l was saved as input sample (1%) at 4°C. Samples were incubated with primary antibodies overnight at 4°C. And then, incubated with 10 *μ*l Dynabeads Protein A for 1 hour at 4°C. After 6 washes with lysis buffer containing 100 mM NaCl and 1% TritonX100, ChIP DNA was eluted twice with 125 *μ*l fresh elution buffer (0.2% SDS, 0.1M NaHCO_3_, 5 mM DTT) at 65°C for 10 min. The input DNA volume was adjusted to 250 *μ*l with elution buffer. To achieve reversal of crosslinking, 1 M Tris-HCl (10 mM final concentration) and 500 mM EDTA (2 mM final concentration) were added to samples. Antibodies used for ChIP were: H3K27me3 (Active Motif, 39155) and H3K4me3 (Millipore 07-473).

#### Library construction

Library construction was performed as previously described (Bowman et al., 2013). In short, after the ChIP sample volume was adjusted to 37.5 *μ*l, end polishing reaction (50 *μ*l) was performed by incubating the sample with 1X T4 ligase buffer (NEB, Ipswich, MA, USA), 0.4 mM dNTPs, 7.5U T4 Polymerase (NEB), 2.5U Klenow polymerase (NEB), 25U polynucleotide kinase for 30 min at 20°C in a thermocycler. To clean-up the samples, Solid Phase Reversible Immobilization (SPRI) beads (Agencourt AMPure XP, Beckman Coulter) were used at a 1.8X beads ratio. Once, DNA was eluted with 16.5 *μ*l water, A-tailing reaction (25 *μ*l) was performed. To do so, 16 *μ*l sample with 1X NEB buffer 2, 0.2 mM dATP, 7.5U Klenow 3’-5’ exo minus (NEB) were incubated for 30 min at 37°C. SPRI cleanup was performed with 1.8X beads ratio and DNA was eluted with 9.5 *μ*l of water. Adapter ligation reaction (25 *μ*l) was performed by incubating 9 *μ*l of sample with 1X rapid T4 ligase buffer (Enzymatics, Beverly, MA, USA), 0.01 uM annealed universal adapter, 150 U T4 rapid ligase (Enzymatics) for 15 min at room temperature. SPRI cleanup was performed once again with 1.6X beads ratio and DNA was eluted with 10.5 *μ*l water.

Finally, library amplification was performed by setting up a PCR reaction (50 *μ*l) with 1X Phusion HF master mix (NEB), 0.2 uM universal primer, 0.2 uM barcoded primer, 1X SYBR Green I (Invitrogen), and 0.5 *μ*l Rox (USB). Then, PCR reaction was performed using an Applied Biosystems 7500 Fast Real-Time PCR System. Program used was as followed: an initial denaturing for 30 seconds at 98°C, followed by multiple cycles of 10 seconds denaturation at 98°C, 20 seconds annealing at 64°C, and 45 seconds extension at 72°C. Reactions were terminated at the end of the extension phase, after SYBR green reported reaction kinetics in the log phase for several cycles.

### Bioinformatics

Reads are aligned to dm3 with bowtie2 (v2.2.4) (Langmead & Salzberg, 2012). Coverage tracks are produced with deeptools2 (v2.5.0.1) (Ramirez et al., 2016) by subtracting the respective input (--ratio subract --normalizeTo1x 121400000 -bs 1). Reads of ChIP alignments are counted with multiBamCov of bedtools (v2.25.0) (Quinlan & Hall, 2010). H3K4me3 reads are counted in a 500bp region downstream of the first TSS. H3K27me3 reads are counted over the genebody. Flybase 5.44 is used as annotation. Differential regions are called with DESeq2 (v1.22.2) (Love, Huber, & Anders, 2014). Heatmaps of differential regions are generated with ComplexHeatmaps (v2.1.0) (Gu, Eils, & Schlesner, 2016). The hierarchical tree is based on log2FC (DEseq2) with method complete and euclidean distance. In addition, log2TPMs are shown.

### Accession numbers

The Gene Expression Omnibus accession number for the ChIP-sequencing data reported in this paper is GSE134509.

### Enrichment analysis

Gene Ontology (GO) enrichment analysis were performed on www.flymine.org/ with Holm-Bonferroni correction with max p-value 0.05. For analysis of protein complexes the Compleat website (https://www.flyrnai.org/compleat/) was used (Vinayagam et al., 2013).

## Acknowledgements

We thank all Knoblich lab members for support and discussions, Francois Bonnay and Joshua A. Bagley for comments on the manuscript, Elke Kleiner, the IMP/IMBA Biooptics Facility for assistance and the Harvard TRiP collection, the Bloomington Drosophila stock center and the Vienna Drosophila Resource Center (VDRC) for reagents.

## Author contributions

M.D.A. and L.L. conducted experiments, interpreted data and wrote the manuscript under the supervision of J.A.K. E.E. optimized the chromatin isolation with S.K.B. who performed ChIP-Seq and under the supervision of R.E.K. T.B. performed the analysis on the ChIP-Seq data.

## Competing interests

The authors declare no competing financial interests.

## Funding

Work in the Knoblich laboratory is supported by the Austrian Academy of Sciences, the Austrian Science Fund (Z_153_B09), and an advanced grant from the European Research Council (ERC).

